# Disease driven extinction in the wild of the Kihansi spray toad (*Nectophrynoides asperginis*)

**DOI:** 10.1101/677971

**Authors:** C. Weldon, A. Channing, G. Misinzo, A.A. Cunningham

## Abstract

The Kihansi spray toad, *Nectophrynoides asperginis*, became extinct in the wild despite population monitoring and conservation management of its habitat in the Kihansi gorge, Tanzania. Anecdotal evidence has indicated human induced habitat modification, predators, pesticides and disease as possible causes of a rapid population decline and the species extirpation. Here, we systematically investigate the role of disease in the extinction event of the wild toad population. The amphibian chytrid fungus, *Batrachochytrium dendrobatidis*, was detected in spray toads that died during the extinction event and subsequently in other amphibian species in Kihansi Gorge and the adjacent Udagaji Gorge, but not in any toads collected prior to this. Following the population decline, the remnant spray toad population gradually disappeared over a nine-month period. We demonstrate how demographic and behavioral attributes predisposed the spray toads to chytridiomycosis, due to *B. dendrobatidis* infection, and how epidemic disease could have been exacerbated by altered environmental conditions in the spray wetlands. Our results show that chytridiomycosis was the proximate cause of extinction in the wild of *N. asperginis*. This represents the first known case of extinction by disease of an amphibian species in Africa. A captive breeding program in the US and Tanzania ensures the survival of the species and a reintroduction program is underway. However, we caution that chytridiomycosis remains an existing threat that requires a comprehensive mitigation strategy before the desired conservation outcome of an established population of repatriated toads can be achieved.

## Introduction

The Kihansi spray toad, *Nectophrynoides asperginis*, was discovered in 1996 at Kihansi Gorge in the southern Udzungwa Mountains, Tanzania and was distinctive in Africa in terms of the habitat it occupied (Poynton et al. 1998). Its entire known distribution was restricted to less than 0.15 km^2^ of a unique vegetation type within a narrow strip of moist forest bordering the Kihansi River in the Kihansi gorge. Continuous spray generated by the Kihansi River as it flowed over the steep Udzungwa scarp previously showered patches of herbaceous vegetation and moss-covered rocks to create a series of spray wetlands. While *N. asperginis* could be classified as naturally rare at the time of its discovery due to its small geographic range (Rabinowitz et al. 1986), within its limited habitable range *N. asperginis* was locally abundant, owing to its characteristically dense populations (Channing et al. 2006).

The spray wetlands underwent severe alteration because of a ten-fold reduction in water flow following the construction of a hydroelectric power plant that was commissioned in May 2000. The bypass flow released from the newly constructed dam above the main Kihansi falls was not sufficient to generate the spray that characterised the gorge and sustain the unique ecosystem within it (Quinn et al. 2005). Immediately following the diversion of the river, the spray toad population experienced a considerable drop in numbers from its original population estimated at almost 18,000 (Channing et al. 2006). An elaborate sprinkler system was installed in three of the five wetlands (Upper, Lower and Mid-Gorge Spray Wetlands) to mitigate the situation by mimicking the spray zone conditions of high relative humidity and low, constant temperatures (NORPLAN 2001). Spray toad numbers continued to fluctuate, but substantially increased from below 2,000 in March 2001 to almost 18,000 in early June 2003 for the Upper Spray Wetland (Channing et al. 2006). In the weeks following the June 2003 census, however, population numbers unexpectedly plummeted and *N. asperginis* was declared extinct in the wild after repeated subsequent surveys yielded no records of the spray toad (IUCN SSC Amphibian Specialist Group, 2015). A captive assurance population established in December 2000, following the initial population decline, prevented the extinction of the species. The offspring of these founder animals were housed at the New York Bronx and Toledo zoos in the U.S.A. (Lee et al. 2006), and, from August 2010 to date, additionally at captive breeding facilities at the University of Dar es Salaam and Kihansi, Tanzania.

The cause of the decline and subsequent extinction in the wild of *N. asperginis* has been widely debated, but little evidence to support any of the proposed causes has been presented (Weldon and du Preez 2004, Quinn et al. 2005, Channing et al. 2006, Krajik 2006). There is wide support that drying of the habitat as a direct result of commissioning of the Lower Kihansi Hydropower Project played a central role in the initial decline of the species. In addition to the obvious habitat modification caused by the reduction in bypass flow (gradual desertification accompanied by alteration of vegetation composition), sediment laden water and pesticide release that coincided with flushing of the dam just prior to the population crash have also been suggested as contributory factors (Hawkes et al. 2008, Rija et al. 2010).

This sudden, catastrophic population crash lasted less than one month and left only a small remnant of the population alive. This remnant population gradually declined over the next eight months until no more toads could be found (Hawkes et al. 2008). A crash of this magnitude is characteristically caused by a stochastic event that pushes a population beyond its resilience threshold. The variation in a number of demographic traits (demographic stochasticity) is proportionally larger, the smaller the population size (Shaffer 1981), but the spray toad population was at its peak in the months preceding the crash and had survived all former fluctuations in population size despite a modified habitat. Similarly, environmental stochasticity is more prone to cause the extinction of small populations. Although the normal range of variation in physical factors was dramatically altered by a temporary shut-down of the sprinkler system during flushing of the dam, the same procedure had been successfully conducted before without adversely affecting the toads. It appears more likely that a catastrophe (e.g. hurricane, major fire, epizootic disease) to which small populations are particularly vulnerable caused the *N. asperginis* population crash.

Interestingly, the fungal pathogen, *Batrachochytrium dendrobatidis*, was detected in some spray toads at the time of the ultimate population crash, thus disease has been proposed as a possible cause of the population decline and extirpation (Weldon and du Preez 2004). The presence of *B. dendrobatidis* in a species that went extinct is noteworthy, since this fungus has been identified as an emerging amphibian pathogen, capable of acting as both the proximate and predisposing cause of amphibian species extinction (Schloegel et al. 2006, Fisher et al. 2009). Assessing the role of pathogens in population declines is challenging because of the logistical and technical difficulties involved (Daszak et al. 2003). Linking infectious disease to extinction requires collection of population data from the last remnant group of a species prior to extinction and pathological examination of at least some of these individuals (MacPhee and Marx, 1997). Here, we investigate the occurrence of the chytrid fungus *B. dendrobatidis* in the Kihansi Gorge region through systematic observations and retrospective surveys and we specifically examine the role of chytridiomycosis as a possible cause of the population crash that led to the extinction of *N. asperginis* in the wild.

## Material and Methods

Kihansi Gorge is situated in the southern Udzungwa Mountains, Tanzania (approximate co-ordinates −8.58333, 35.8500) and holds a narrow strip of rain forest surrounded by Miombo woodland. A unique vegetation type grows within the spray zone of the Kihansi River waterfalls to form a series of spray wetlands (Figure 1). The adjacent Udagaji Gorge (approximate co-ordinates −8.58666, 35.87333) was selected as a reference site because it does not share the same catchment as the Kihansi River, but has a similar amphibian community structure as Kihansi Gorge, apart from *N. asperginis*.

**Figure 1.**
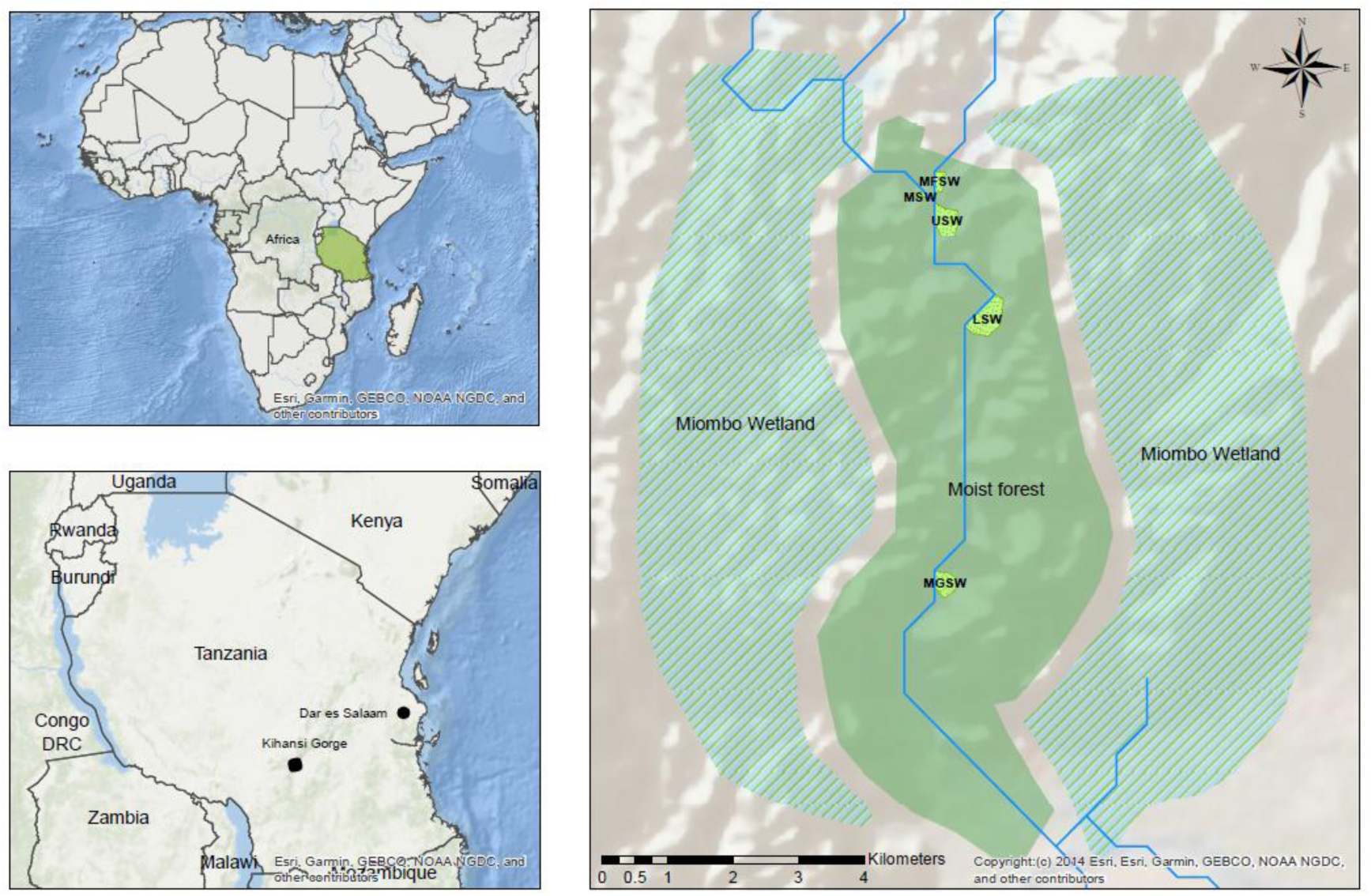
Map of Kihansi Gorge indicating the course of the Kihansi River and location of spray wetlands. MFSW, Main Falls Spray wetland; MSW, Mhalala Spray wetland; UPS, Upper Spray wetland; LSW, Lower Spray wetland; MGSW, Mid-gorge Spray wetland.

### Timeline of decline and retrospective survey

Observations from the sprinkler-service and monitoring team at Kihansi, which were mostly disclosed in the report of a Population and Habitat Viability Assessment for the spray toads (Lee et al. 2006, Hawkes et al. 2008), were examined to compile a timeline of the estimated size of the remnant *N. asperginis* population until extinction.

Using inter-dental brushes (3.2 to 6.0 mm; Oral B Laboratories), skin brushings of the ventral abdomen, hind limbs and hind feet were taken of the paratype collection of *N. asperginis* at the Natural History Museum, London (see Poynton et al. 1998) as described by Soto-Azat et al. (2009). DNA was extracted from each sample and was tested for the presence of *B. dendrobatidis* using a specific real-time polymerase chain reaction (qPCR) assay as described by Boyle et al. (2004). For each sample, the diagnostic assay was performed in duplicate, and standards of known *B. dendrobatidis* zoospore concentrations and negative controls were included within each qPCR plate. A sample was considered to be positive when: (1) amplification (i.e. a clearly sigmoid curve) occurred in both replicated qPCR reactions, and (2) values > 0.1 genomic equivalents (GE) were obtained from both replicated reactions.

A toe clip was taken from the left hind foot of each archived *N. asperginis* and *N. tornieri* specimens housed in the herpetofauna collection of the University of Dar es Salaam and the University of the Western Cape. These specimens were collected alive during monitoring expeditions, 1997-2002, under the auspices of the Lower Kihansi Environmental Management Project (LKEMP). Each toe clip was stored in 70% alcohol and prepared for histological examination using routine methods. Tissue sections, 6 μm thick, were stained with Mayer’s haematoxylin and counter stained with eosin. Slides were then examined using a Nikon Eclipse E800 compound microscope for the presence of *B. dendrobatidis* thalli using the criteria described by Berger et al. (1999).

In June to August 2003, during the terminal population crash, eight *N. asperginis* found dead in the Upper and Mid-Gorge Spray Wetlands were collected by LKEMP staff. Part of the left hind foot of seven of these carcases was similarly prepared for histological examination. One of the specimens was swabbed with a sterile cotton swab and analysed using conventional PCR to test for the presence of *B. dendrobatidis*, as previously described by Annis et al. (2004).

### Field survey

Field trips to Kihansi and Udagaji gorges were undertaken during November 2003 and May 2006. Before entering and upon exiting a gorge or spray wetland, footwear was scrubbed and disinfected with a 2% sodium hypochlorite solution. Kihansi Gorge was thoroughly searched for the presence of amphibians both during the day and at night, concentrating on the spray wetlands and immediate surrounding forest. Diurnal surveys involved systematically searching among wetland vegetation and exposed rock surfaces inside wetlands, using the sprinkler lines as transects. Nocturnal surveys involved searching among wetland vegetation along the artificial walkways and examining all exposed rock surfaces. Frogs and toads were collected by hand and kept individually in separate plastic bags. Disposable nitrile gloves were worn whenever animals were handled and a clean pair was used for each animal. The fifth toe of the left hind foot of each animal captured was surgically removed and fixed in 70% alcohol for later examination using histology. The wound was anointed with antiseptic ointment (Betadine; Adcock Ingram Ltd.) before the animal was released at the point of capture. Scissors were sterilized with alcohol wipes between each sample. At Udagaji Gorge, searches for amphibians took place along the banks of the river (5 m on either side) up to the main falls of the gorge. The same sampling protocol was followed as for Kihansi Gorge.

## Results

### Population trend

By July 2003, spray toad population numbers had plummeted from an estimate 17,745 (4.3 toads/m^2^) for the upper spray wetland in June 2003 to only 43 toads in total for all wetlands combined. Converting all subsequent records of toads to numbers per search hour revealed that a remnant population persisted after the population crash for nine months. During the first four months numbers continued to drop sharply from 31 toads/h in July 2003 to 3.2 toads/h in November 2003, after which it remained more or less stable between 2.5 and 1.3 toads/h (Figure 1) until March 2004 when the last confirmed sighting of the Kihansi spray toads in the wild was made.

### Retrospective survey

A total of 107 archived *N. asperginis* and eight *N. tornieri* from Kihansi Gorge were tested for chytrid fungus infection using histological examination of toe clips. All of the examined specimens were collected from the wild during 1996 or 2003 and included the 18 paratypes used in the original species description of *N. asperginis*. No archived amphibians from Udagaji Gorge were available for disease screening.

All 99 *N. asperginis* as well as the eight *N. tornieri* that were collected between 1996 and 2002 tested negative for chytrid fungus infection (Table 1). However, five of the eight Kihansi spray toads found dead in 2003 (Upper and Mid-Gorge Spray wetlands) had chytridiomycosis (Tables 1 & 2) due to infection with chytrid fungal zoosporangia which were morphologically consistent with those of *Batrachochytrium dendrobatidis* (Berger et al. 1998, 1999), and one tested positive for *B. dendrobatidis* with PCR. Histopathological examination of these toads consistently revealed severe infection intensity concentrated in the anterior half of the feet (Figure 2). Almost 50% of the examined skin surface contained chytrid thalli, with infected loci often consisting of up to 10 layers of sporangia.

**Table 1.**
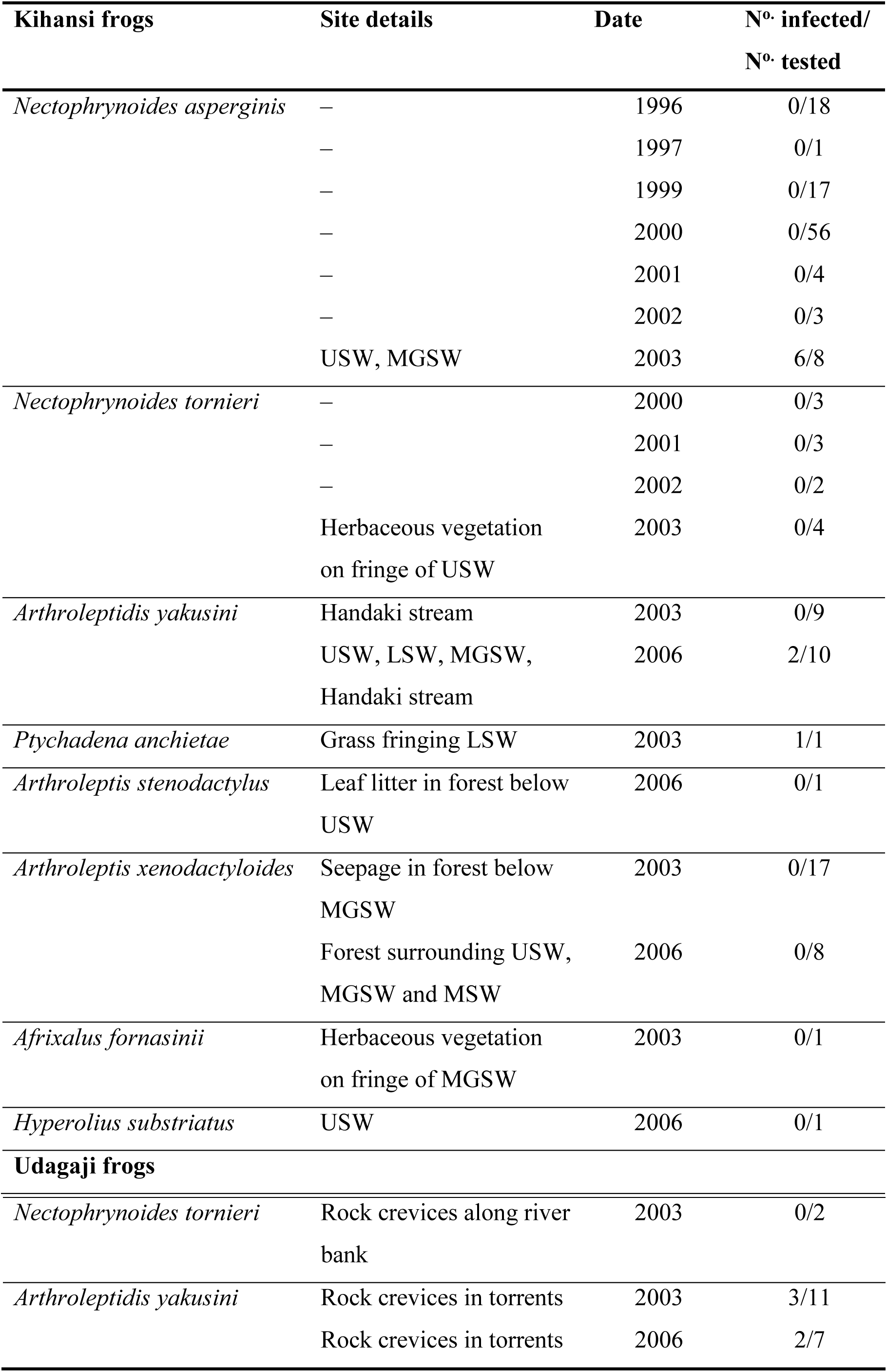
Occurrence of chytrid fungal infection in amphibians from Kihansi and Udagaji gorges since the discovery of *Nectophrynoides asperginis*. USW – Upper Spray Wetland, MGSW – Mid-Gorge Spray Wetland, LSW – Lower Spray Wetland, MSW – Mhalala Spray Wetland

**Table 2.** *Nectophrynoides asperginis* infection history since the earliest detection of chytrid fungus, which coincides with the timing of the population crash. Histology was used in most diagnosis, except for one specimen (asterisk) for which PCR was used.

| Date | Nº. specimens | Nº. infected | Locality |
| --- | --- | --- | --- |
| 4 Jun 2003 | 2 | 1 | Upper Spray Wetland |
| 28 Jul 2003 | 3 | 2 | Mid-Gorge Spray Wetland |
| 4 Aug 2003 | 1 | 1* | Mid-Gorge Spray Wetland |
| 14 Aug 2003 | 2 | 2 | Mid-Gorge Spray Wetland |

**Figure 2.**
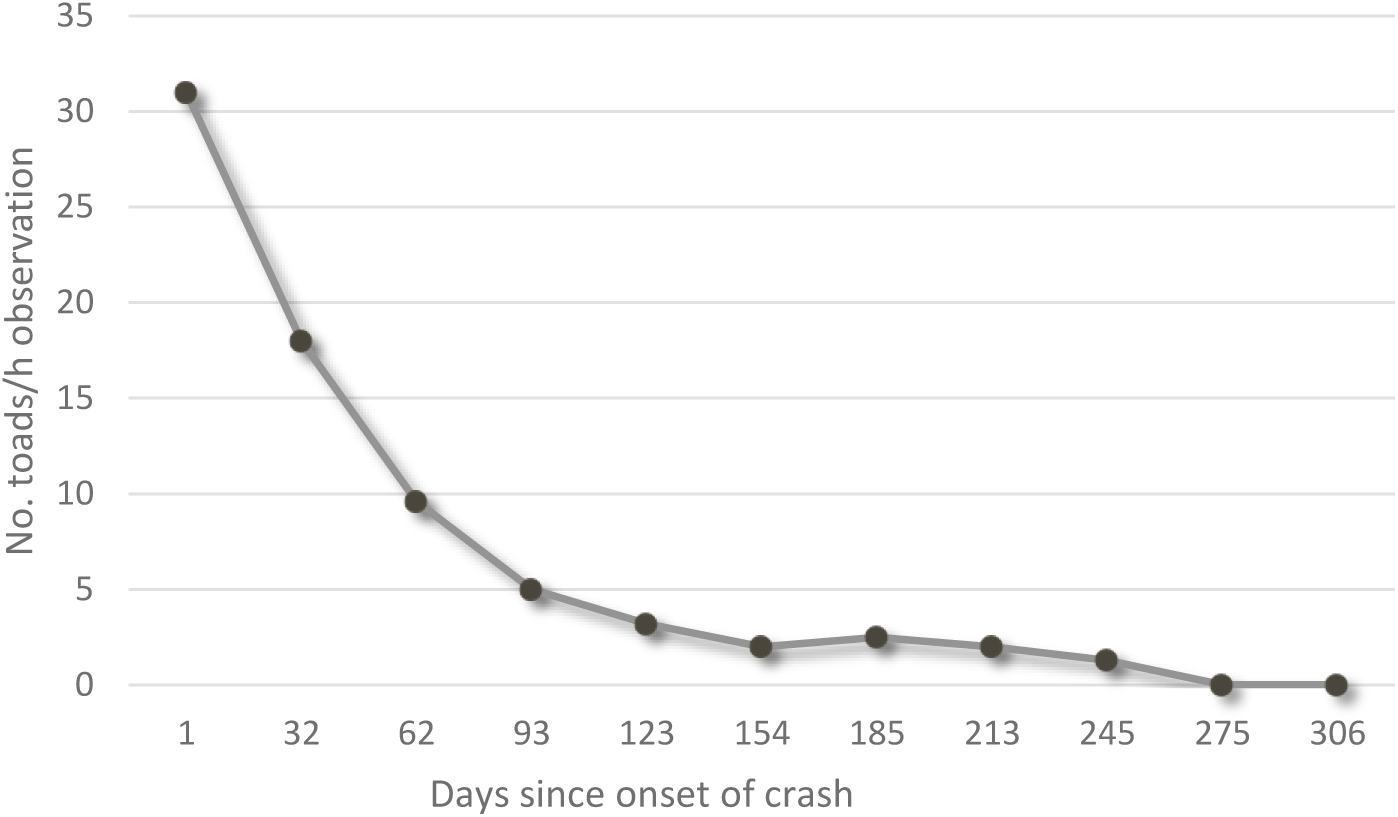
Population trend of *Nectophrynoides asperginis* following the crash in June 2003. Numbers are based on total number of toads observed per hour of search effort (compiled from Hawkes et al. 2008).

**Figure 3.**
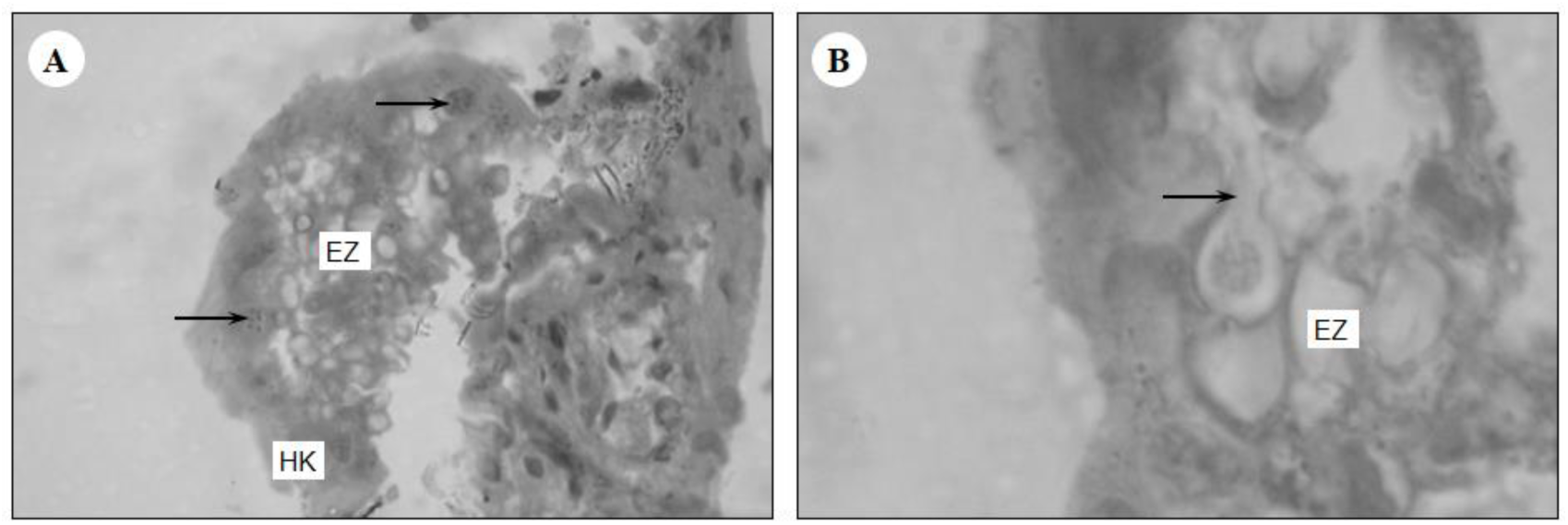
Micrographs of hematoxylin and eosin stained sections of *Nectophrynoides asperginis* skin. A) A large cluster of mostly empty zoosporangia (EZ) in a fragment of partially detached epidermis, with severe hyperkeratosis in the *stratum corneum* (HK). Zoosporangia containing zoospores (arrows) are also visible. B) Zoosporangium with discharge papilla (arrow), surrounded by empty zoosporangia within the *stratum corneum* with advanced autolysis.

### Field survey

The field survey included the screening of 60 specimens from seven amphibian species from Kihansi Gorge, and 20 specimens from Udagaji Gorge (Table 1). Two species from Kihansi Gorge tested positive for infection with chytrid fungus, morphologically consistent with *B. dendrobatidis*, on histological examination of toe-clips, namely *Ptychadena anchietae* and *Arthroleptides yakusini* (Table 1). Infection was also detected in *A. yakusini* from Udagaji Gorge in 2003 and 2006 (27.3% and 28.6% prevalence respectively).

Despite intensive diurnal and nocturnal surveys (32 man-hours in 2003, 21 man-hours in 2006), no Kihansi spray toads could be found in any of the spray wetlands where the species was known to have occurred. In both years, the spray wetlands were almost totally devoid of any anurans, but for a few *A. yakusini* and a single *Hyperolius substriatus* in 2006.

## Discussion

### Evidence linking chytridiomycosis with the extinction of Kihansi spray toads in the wild

The rapidity and severity with which the spray toad population declined is typical of an infectious disease epidemic in a naïve host (Rachowicz et al. 2005).

Chytridiomycosis is known to have had such a devastating effect on amphibian populations elsewhere (Hudson et al. 2016), and the pattern of decline in *N. asperginis* shows similarity to amphibian declines attributable to chytridiomycosis elsewhere (Lips et al. 2006, Hudson et al. 2016).

According to Schloegel et al. (2006), extinction can be attributed to infection if a pathogen caused declines in the majority of a species’ population leading to its extinction; if a pathogen was responsible for the die-off of the last remnant population of a species regardless of the cause of previous declines; or if a pathogen caused the death of the last individual of a species even if other factors were responsible for driving the species to the verge of extinction. The evidence presented here indicates that *B. dendrobatidis* infection caused the decline of *N. asperginis*, eventually leading to the species becoming extinct in the wild: (1) The *N. asperginis* population underwent a rapid and catastrophic decline due to adult mortality. (2) Chytridiomycosis was prevalent among animals examined that had been found dead during this population crash. (3) Histopathological examination of the skin of Kihansi spray toads that died during the population decline demonstrated an acute infection with severe and extensive epidermal lesions associated with zoosporangia morphologically consistent with *B. dendrobatidis*, and *B. dendrobatidis* infection was confirmed using PCR in one of the last specimens of the species to be collected before its extinction. (4) None of the amphibians tested that had been collected over six years prior to the population crash were positive for *B. dendrobatidis*. The epidemiological chain of events, therefore, indicates that the timing of the population crash coincided with the first appearance of chytridiomycosis in *N. asperginis* as well as in the related *N. tornieri*. Based on this evidence we are assuming that all animals that died with chytridiomycosis died from *B. dendrobatidis* infection.

The infection dynamics of *B. dendrobatidis* during frog die-offs is typically characterized by a rapid buildup of high-level infections, followed by death due to chytridiomycosis (Vrendenburg et al. 2010). The first arrival of *B. dendrobatidis* into a naive frog population often coincides with an outbreak of chytridiomycosis (Berger et al. 1998, Lips et al. 2006, Vrendenburg et al. 2010, Hudson et al. 2016). A similar epidemiological pattern to Kihansi Gorge (no evidence of infection until just before the observed frog die-offs) was consistently witnessed at multiple sites in California’s Sierra Nevada that resulted in the extirpation of numerous mountain yellow-legged frog (*Rana muscosa*) populations (Vrendenburg et al. 2010). This pattern was also found with the mountain chicken frog (*Leptodactylus fallax*) decline on the Caribbean islands of Montserrat and Dominica (Hudson et al. 2016). Similarly, pathological examinations on declining populations of the now extinct sharp-snouted torrent frog (*Taudactylus acutirostris*) from Australia, including individuals from the last free-living remnant population, demonstrated chytridiomycosis as the probable cause of extinction of this species (Schloegel et al. 2006). Since not a single animal from the remnant *N. asperginis* population was collected during the nine months of surveying following the crash, *B. dendrobatidis* could not be confirmed as having caused the deaths of the last few animals that lingered on as a remnant group, but whether or not they were killed by *B. dendrobatidis*, according to the criteria of Schloegel et al. (2006), there is compelling evidence that this pathogen caused the extinction of *N. asperginis*.

The absence of *B. dendrobatidis* from all historic records predating the population crash does not support the notion of a chytridiomycosis outbreak that emerged from an endemic infection triggered by changing environmental conditions (chytrid thermal optimum hypothesis) (Pounds et al. 2006). Rather, the emergence of chytridiomycosis at Kihansi Gorge suggests that *B. dendrobatidis* had been introduced to a naïve *N. asperginis* population shortly before the epidemic occurred (novel pathogen hypothesis) (Lips et al. 2008, James et al. 2009). Moreover, the absence of infection in any of the founder toads that were translocated to the United States in 2000 (Lee et al. 2006), provides further evidence that *B. dendrobatidis* had been introduced to the wild toad population shortly before the terminal population decline.

### Predisposition of Kihansi spray toads to chytridiomycosis

Species extinction represents one end of a spectrum of possible outcomes of infection with *B. dendrobatidis*. In the most benign cases, individuals of some amphibian species carry the infection with apparently little to no effects (Daszak et al. 2004, Weldon et al. 2004), whereas other species experience death of individuals, extirpation of populations or species extinctions (Skerrat et al. 2007, Scheele et al. 2019). Previous studies have demonstrated that a combination of physiological, demographic, and ecological traits at the species level (e.g. Woodhams et al. 2006, Smith et al. 2009, Briggs et al. 2010), as well as the virulence of the particular strain of *B. dendrobatidis* (Ferrar et al. 2011, O’Hanlon et al. 2018) influence the outcome of infection.

In general, species that are rare, that have low fecundity, that have aquatic larvae associated with streams, or that occur at high elevation are predisposed to population declines and extinction due to *B. dendrobatidis* infection (Lips et al. 2003, Murray and Hose 2005, Bielby et al. 2008). Except for aquatic larvae, *N. asperginis* displays all of these characteristics (600–940 m alt., home range < 2 ha, prolonged gestation, live bearing, clutch size of 5–16; Poynton et al. 1998, Channing et al. 2006), suggesting that it was at high risk of population decline due to chytridiomycosis. The exceptionally high densities at which spray toads used to occur (up to 17 toads/m^2^), together with the habit of congregating on exposed rocks within wetlands (Poynton et al. 1998), implies that substantial physical contact occurred between individuals. It has been demonstrated that *B. dendrobatidis* transmission is density dependent and that host population density governs host-pathogen dynamics (Briggs et al. 2010). Thus it is likely that the population density and behaviour of *N. asperginis* facilitated frog-to-frog transmission and enabled the rapid spread of *B. dendrobatidis*.

### Environmental exacerbation of the epidemic

*Nectophrynoides asperginis* appears to be susceptible to lethal chytridiomycosis due to *B. dendrobatidis* infection in the absence of any predisposing factors. In addition to the epidemic mortality seen in the wild, all of the spray toads in a single enclosure in the Wildlife Conservation Society’s Bronx Zoo died from a chytridiomycosis outbreak following an incursion of *B. dendrobatidis* (McAloose et al. 2008). The species’ high susceptibility to chytridiomycosis was further demonstrated when 62% of the captive breeding facility population in Kihansi died of the disease in under seven weeks following incursion of *B. dendrobatidis* despite daily removal of carcases and antifungal treatment (Makange et al. 2014).

In addition to the species’ susceptibility to lethal chytridiomycosis, the prevailing environmental conditions in the spray wetlands as well as anthropogenic disturbance of these conditions just prior to the 2003 population crash, might have exacerbated epidemic chytridiomycosis. A constant relative humidity of near saturation and ambient temperature of 15–23°C (Poynton et al. 1998) provides optimal growth conditions for *B. dendrobatidis* (see Longcore et al. 1999). These favourable conditions would have facilitated the rapid invasion of the spray wetlands following incursion by the fungus in the presence of a suitable host. Furthermore, the constant movement of saturated air generated by the spray zones might have aided the dissemination of *B. dendrobatidis* between adjacent wetlands. Shortly before the disease outbreak occurred in the wild, there was a temporary shutdown of the sprinklers during intermittent high-flow tests of the dam. Spray toads reacted to such periods of reduced humidity by aggregating in parts of the wetland closest to the waterfall that still received small amounts of natural spray (Channing et al. 2006). The aggregation of amphibians in damp refugia during periods of drought increases the rate of *B. dendrobatidis* transmission and has been documented elsewhere as a trigger for chytridiomycosis outbreaks (Burrowes et al. 2004, Longo et al. 2010). The ultra-dense assemblages of spray toads during the June 2003 high-flow test also would have accelerated frog-to-frog disease transmission.

### Implications for conservation

A reintroduction plan for *N. asperginis* was developed in 2010 that involved maintaining captive assurance colonies at the University of Dar es Salaam and the Kihansi Research Centre, and the reinstatement of a viable wild population at Kihansi Gorge (Rija et al. 2010). In addition to identifying a feasible chytridiomycosis mitigation strategy, the successful establishment of a wild spray toad population will depend on overcoming other management challenges such as water flow and quality, vegetation change and the encroachment of non-native species (Rija et al. 2010).

Our results show that *B. dendrobatidis* is likely to persist in Kihansi Gorge in other amphibian species; it was detected even more recently in *Phrynobatrachus mababiensis* from below the gorge (Makange et al. 2014). Sympatric amphibian species can act as reservoir hosts of *B. dendrobatidis* and transmit the pathogen to any repatriated spray toads. Continued monitoring and assessment of the risks of disease should be a key component of the reintroduction strategy for any species. This is particularly important for *N. asperginis* because the extinction of this species in the wild was not observed until it was imminent; this, despite monitoring and conservation strategies to maximize the preservation of the species at that time. Garner et al. (2016) highlighted the need for field trials to develop chytridiomycosis mitigation in the wild, but also recommended that such *in situ* work be guided by the results of *ex situ* research. The current captive breeding and reintroduction program for *N. asperginis* provides a fitting opportunity for doing this as a conservation action for this species, but also for furthering the science of infectious diseases mitigation for threatened wildlife.

## Conclusion

Our investigation into the rapid, catastrophic decline of *N. asperginis* suggests that the virulent amphibian pathogen, *B. dendrobatidis*, was responsible for driving this toad to extinction in the wild. This represents the first documented case of extinction by disease in the wild of an amphibian species in Africa. The extremely restricted distribution of the species made it vulnerable to a stochastic event such as infectious disease introduction and this was enhanced by the high population density, the aggregation behaviour of the toads and the optimal environmental conditions for *B. dendrobatidis*. The live bearing life history of the species added to its vulnerability because of its low fecundity and prolonged gestation. There is a lack of evidence of the cause of extinction of the remnant Kihansi spray toads that survived the chytridiomycosis epidemic, but as the population declined and population density decreased, infection dynamics dictate that a small number of animals survive through the tail of the epidemic curve. Also, the epidemic likely left surviving *N. asperginis* with population numbers below a threshold number of individuals required to maintain the characteristic behaviour associated with a species that forms large aggregations. Whether these remaining animals died of *B. dendrobatidis* infection transmitted from other Kihansi spray toads, sympatric amphibians or due to other factors, we posit that this pathogen was the cause of this species’ extinction in the wild.

## Acknowledgements

This research was funded by the Lower Kihansi Environmental Management Project (LKEMP), who also provided logistical support and assistance during fieldwork. We thank Kim Howell and David Moyer for collecting many of the archived spray toads that were included in this study. John Wood (Pisces Molecular) conducted PCR diagnostic tests on one of the specimens. Celia Cloete developed the map of Kihansi Gorge.

